# Interacting fruit flies synchronize behavior

**DOI:** 10.1101/545483

**Authors:** Ugne Klibaite, Joshua W. Shaevitz

## Abstract

Social behaviors are ubiquitous and crucial to an animal’s survival and success. The behaviors an animal performs in a social setting are affected by internal factors, inputs from the environment, and interaction with others. To quantify social behaviors, we need to measure both the stochastic nature of behavior of isolated individuals and how these behaviors change as a function of the environment and features of social interaction. We probed the behavior of male and female fruit flies in a circular arena as individuals and within all possible pairings. By combining measurements of the animals’ position in the arena with an unsupervised analysis of their behaviors, we fully define the effects of position in the environment and the presence of a partner on locomotion, grooming, singing, and other behaviors that make up an animal’s repertoire. We find that geometric context tunes behavioral preference, pairs of animals synchronize their behavioral preferences across trials, and paired individuals display signatures of behavioral mimicry.

## Introduction

Social behaviors are exhibited by a wide variety of species and include such diverse categories as courtship, aggression, dominance hierarchies, collective flocking, and group decision making (*Lorenz, 2002*; *Tinbergen, 1963*; *Bialek et al., 2014*; *Giuggioli et al., 2013*; *Ni and Ouellette, 2015*; *Durisko et al., 2014*; *Louis and de Polavieja, 2017*; *Dombrovski et al., 2017*; *Ramdya et al., 2017*). Ultimately, the actions an animal performs are influenced by a combination of environment, social interaction, and internal state (*Censi et al., 2013*). Separating out these contributions to isolate the nature of social interaction has remained a challenge. Many previous attempts to quantify social interactions and networks have used proximity data but this is insufficient to probe social effects on non-locomotive behaviors (*Katz et al., 2011*; *Herbert-Read et al., 2011*). Other work has examined the effect of unidirectional social stimulus on an individual which does not allow for the study of the inherent closed-loop, multi-directional nature of many social behaviors (*Coen and Murthy, 2016*; *Coen et al., 2016*; *Stowers et al., 2017*). We move beyond these limitations by recording the position and full behavioral repertoire of pairs of fruit flies, allowing us to separately quantify the effects of environment and social interactions on behavior.

In pairs from various species, both individuals can perform the same behavior simultaneously, a phenomena referred to as mimicry, imitation, synchrony, and contagion *Zentall* (*2004*, 2006). This is not to be confused the Batesian mimicry, where one species copies the physical appearance of another *Zentall* (*2006*). Neural mechanisms of behavioral mimicry have been explored in humans and the discovery of mirror neurons, cells which reliably fire during the execution of motor sequences performed by others, has spurred hypotheses about their mechanisms in learning and pro-social behavior *Iacoboni* (*2009*). The proposed functional roles of mirror neurons in humans and certain apes include action understanding and action learning, properties which are not likely to exist in simpler organisms *Rizzolatti and Craighero* (*2004*). Behavioral mimicry can result in benefits to both individuals such as an enhanced avoidance of a predator and the facilitation of feeding behaviors *Welty* (*1934*); *Allee et al.* (*1931*); *Tolman* (*1964*); *Bates and Byrne* (*2010*). These benefits can arise from simple behavioral matching, where an animal performs a behavior already present in its repertoire in response to the action of another, and do not require complex cognitive mechanisms *Byrne* (*2009*).

We ask whether there are discernible behavioral effects arising from simple social pairings of fruit flies by using an unsupervised behavioral quantification paradigm. We compare the effects of same-sex and courtship pairings on behavior and find that the behaviors performed by an individual depend on not only the social pairing but also the particular context an individual finds itself in. We find behavioral effects induced by the location of a fly within the experimental arena and the distance of the individual to its interaction partner, as well as behavioral mimicry within interaction pairs at short time scales.

## Results

We recorded the behavior of *Drosophila melanogaster* across lone and paired individuals in a featureless circular arena of radius *R*_arena_ for 30 minute recordings, or less if copulation was reached in the case of courtship pairings. The number of experiments and total time recorded for each pairing is summarized in 2. Video was recorded from above on a backlit stage, where flies were allowed to freely move and interact under a plastic dome with 22mm diameter. Analysis of the resulting video used an unsupervised behavioral paradigm to assign behavioral labels. We also recorded the position and orientation of each individual over time.

### Position and Orientation for Single and Paired Flies

We started by measuring the position of the animals in the arena in each context. Flies show a preference for the edge compared to the middle of the arena, which we quantify through the distance to the center, *d*_*c*_ (Fig. 1A). This effect has been previously reported when flies are restricted to a circular arena, and shows the importance of arena geometry when considering spontaneous behavior (*Valente et al., 2007*).

**Figure 1.**
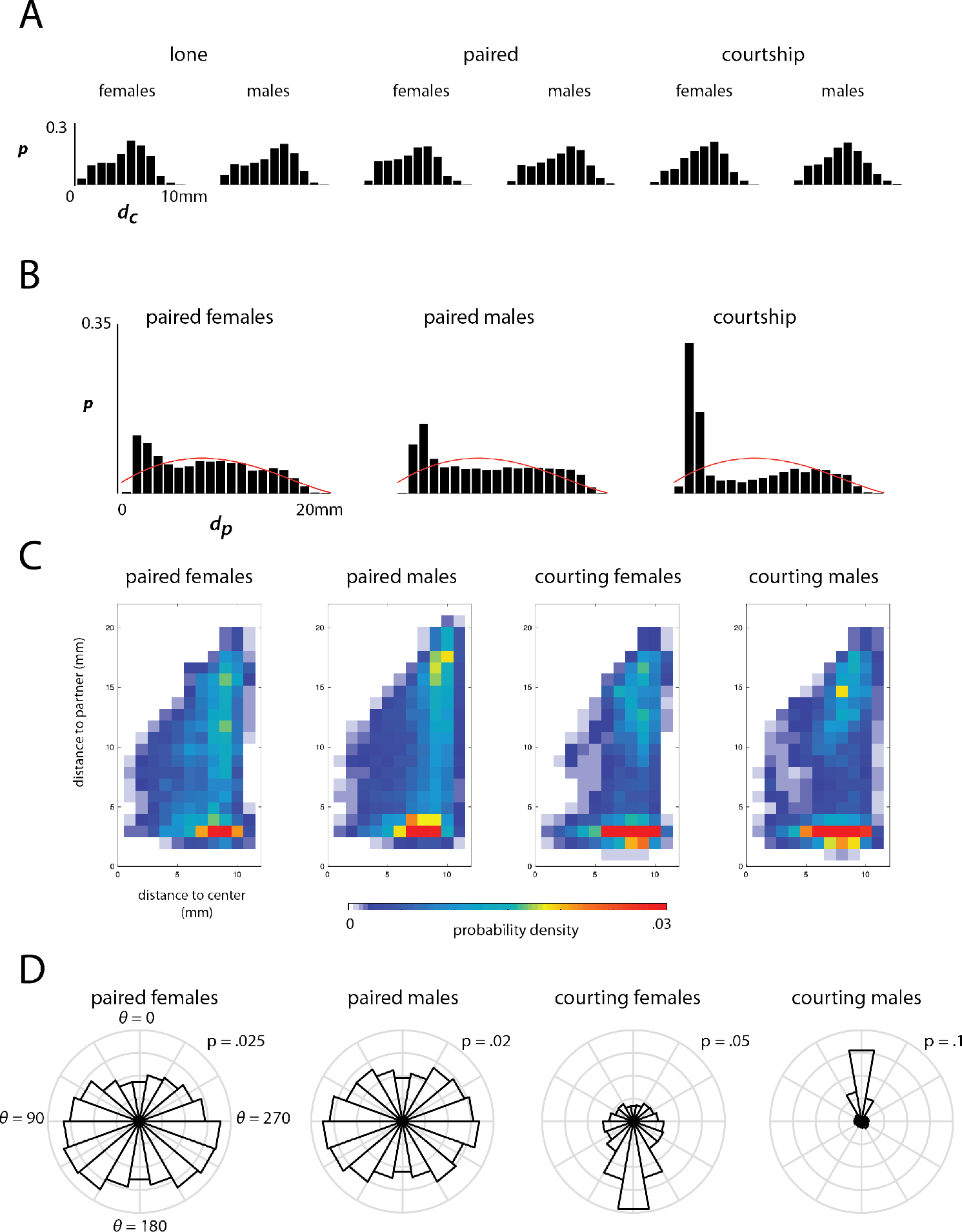
Histograms of distance to the center of the arena, distance to partner, and heading to partner are plotted where applicable. a) The fraction of time individuals from each context reside in ten equal-area concentric circles radiating from the center of the arena outwards. b) Histogram of the distance between paired individuals for all individuals in a given context. The red line indicates the expected distribution of distances between random points within the arena. c) 2-dimensional histogram of residence time in each paired context broken down by distance from the center of the arena (x-axis) and distance to the interaction partner (y-axis). d) Radial histogram of the heading of an individual given a context in relation to the centroid of the arena partner where *θ* = 0 refers to heading directly toward the interaction partner.

For paired contexts, we also measured the distance between the individuals, *d*_*p*_ (Fig. 1B). These distributions are all bi-modal with a narrow peak at short distances and a broader tail that extends to the largest distance *d*_*p*_ = 2*R*_arena_. We compare these histograms to the expected distribution of distances between points on a disk, derived as described in *Weisstein* (*2000*), and find that all pairings spend more time at short distances than expected. We find that individuals in courting pairs spend much more time close to each other than those in same-sex pairs. This is likely because most of the behaviors associated with courtship occur at a short distance to allow for physical, chemical, and auditory communication and attempted copulation. Interestingly, the male-male and female-female pairs also exhibit a peak at short distances, indicating that these pairing also induce short-range social interactions. To investigate the combined effects of pairing and the environment, we plotted the two-dimensional histogram, *P* (*d*_*c*_, *d*_*p*_), for each context (Fig. 1C).

We next examined the prevalence of flies to orient themselves relative to a partner (Fig. 1D). We define the angle of heading for a paired individual as the displacement in degrees from facing toward the interaction partner as described in *Klibaite et al.* (*2017*). We find that the courting male predictably spends most of his time angled toward the female and the female is unlikely to face toward the male during courtship. The same-sex pairings exhibit more uniform distributions of heading, indicating that they do not prefer to face towards or away from their interaction partner as strongly as during courtship. Interestingly, the male individuals show a suppression of heading directly toward their interaction partner. This effect has been suggested previously, when it was shown that a female odor incited males to orient toward and touch other flies, and that a male odor induced the same effect to a lesser extent (*Shorey and Bartell, 1970*).

### Quantification of Behavior Across Social Contexts

We quantified the behaviors exhibited by lone and paired flies using the method described in (*Berman et al., 2014*; *Klibaite et al., 2017*). Briefly, each time point from a movie is mapped to a point in a two-dimensional representation of the postural dynamics of the body (Fig. 2). The estimated two-dimensional probability density, visualized as a heat map such as in Figure 2b, describes the behavioral repertoire of a set of individuals. Clusters from the two-dimensional point cloud, found using a watershed algorithm on the estimated density, represent distinct stereotyped behaviors.

**Figure 2.**
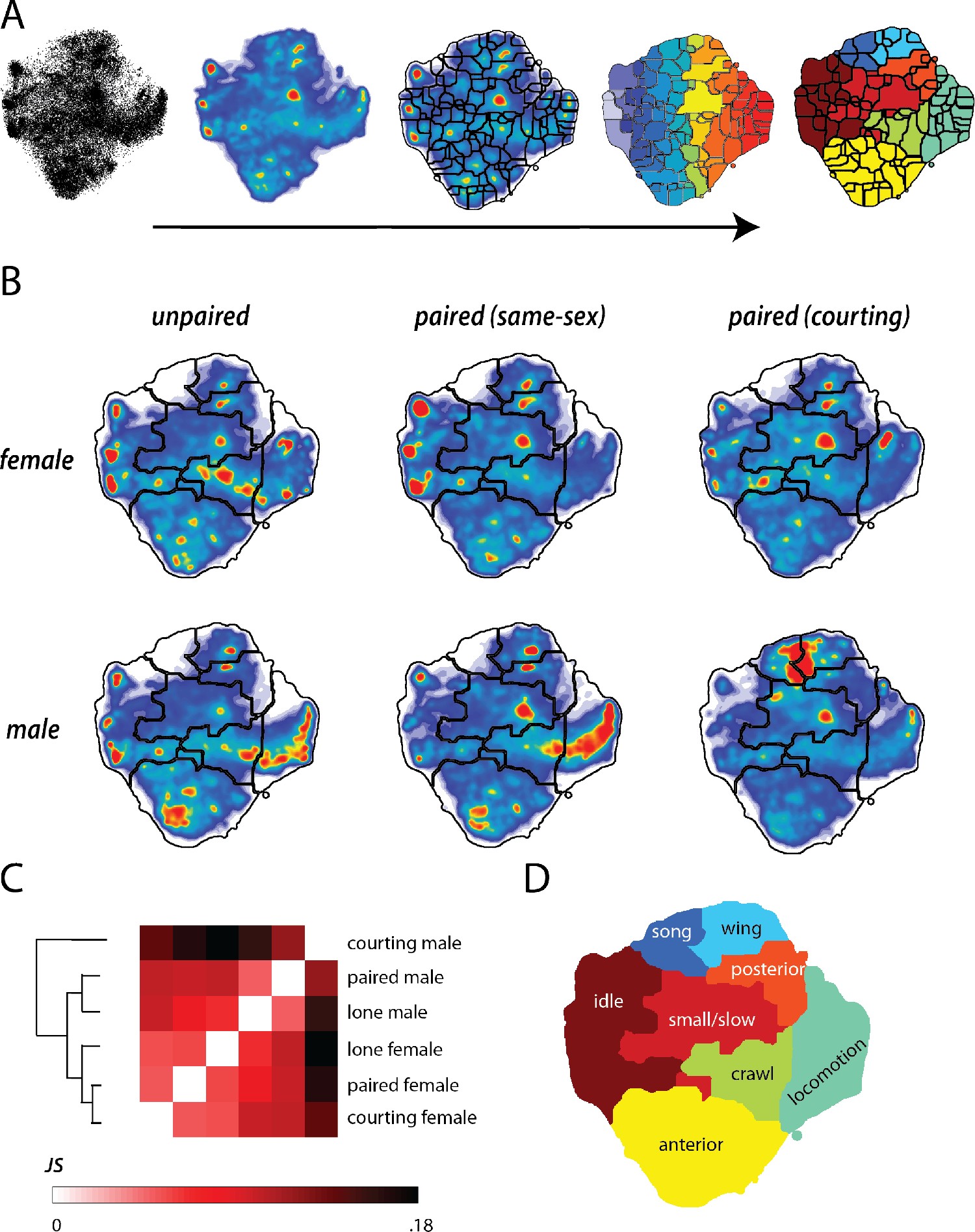
Behavioral densities across paired and lone conditions. a) Representative images from each of five steps used to label behaviors: in sequence the images describe the progression of time-points on the two-dimensional behavioral space through Gaussian filtering, application of the watershed transform, and hand-labeling in order to summarize descriptions of behaviors performed over a long period of time. b) Behavioral densities for each of the six categories of individuals addressed in this publication: lone male and female individuals, same-sex pairs, and male and female courtship pairs. c) We compare the densities across the six conditions by visualizing the JS divergence (Eq. 1) across all groups, and cluster based on similarity. d) Manually assigned labels describe the eight broad categories of behavior exhibited during the behavioral movies and where they fall on the two-dimensional behavioral embedding.

We find that behavior varies based on social context (lone, or paired with a same-sex partner or opposite-sex courtship partner, Fig. 2b). Males perform more fast locomotion behaviors than females when isolated in the experimental chamber, while females tend to crawl and turn more of-ten. Lone individuals of both sexes perform anterior grooming more than their paired counterparts, but paired individuals perform more posterior and wing grooming across all pairings. Females in a same-sex pairing are idle much more often than any individuals in any other experimental group. Individuals in any of the paired contexts display an increase in small body movements (such as reaching using the legs) over isolated individuals. Locomotion decreases from the lone baseline in each paired context except for male-male pairings, where there is a sharp increase. Finally, male individuals in same-sex pairings also display a decrease in idle behaviors over their lone counterparts.

To compare behavior quantitatively, we calculated the Jensen-Shannon (JS) divergence between the behavioral probability densities (*Lin, 1991*, Eq. 1, Fig. 2c). We find that the three female contexts are most similar to each other, followed by the lone and male-male paired contexts. Courting males are by far the most different, driven by courtship-specific song and wing behaviors. Interestingly, males that are lone or paired with another male are more similar to females from all contexts than to courting males.

### The effect of arena position on behavior

We combine measurements of animal position and postural dynamics to investigate the effect of environment on behavior. We started by investigating the behavior of lone individuals at different distances from the center of the arena (Fig. 3). The likelihood of performing wing behaviors diminishes towards the edge of the arena for both sexes, as does the likelihood of being idle. Males are less likely to perform anterior grooming as they reach the edge of the arena, while females exhibit the opposite pattern. Finally, males and females both locomote more when positioned near the edge of the arena, as has been reported previously (*Valente et al., 2007*). Males in particular spend a large portion of time circling the edge of the arena. Overall, the males seem more affected by the placement near the edge, although both sexes equally avoid inhabiting the outer ring of the arena (Fig. 1A).

**Figure 3.**
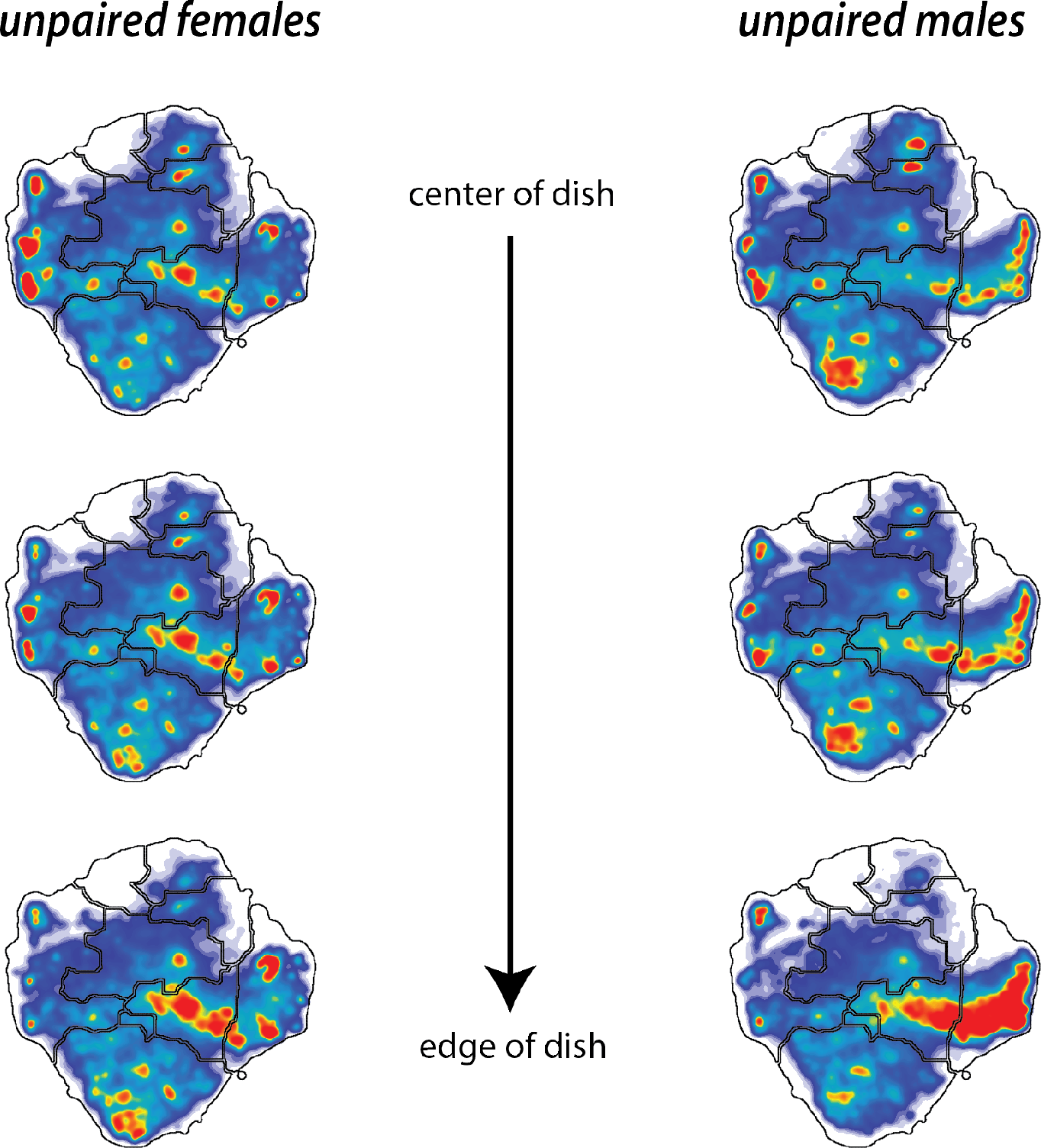
Behavioral density maps of lone individuals corresponding to location in the experimental arena. Density maps generated by embedding behaviors exhibited only when the fly resides in each of three equal-area concentric circles that tile the experimental arena (centered at R=3.15, 7.66, 9.99mm where *R*_arena_ = 11*mm*). Female and male densities are shown separately and reveal that flies are more likely to perform locomotion and much less likely to groom when near the edge of the recording arena, as opposed to when occupying space toward the center of the arena.

Paired flies show the same trend as a function of distance, and we see enriched locomotion at the edges of the arena (Fig. 4). Behavior does not depend linearly on distance from the center of the arena. We quantified the amount of change in behavior with position by computing the JS divergence between behavioral densities at different radii (Fig. 5 A). For all contexts, maps generated from time points where flies are occupying bins within 7*mm* of the center are fairly similar whereas there is a marked change in behavior at the 7*mm* mark. This effect is stronger in males than females, and is not as prominent in the courtship condition for either sex, likely due to social arousal interfering with environmental cues.

**Figure 4.**
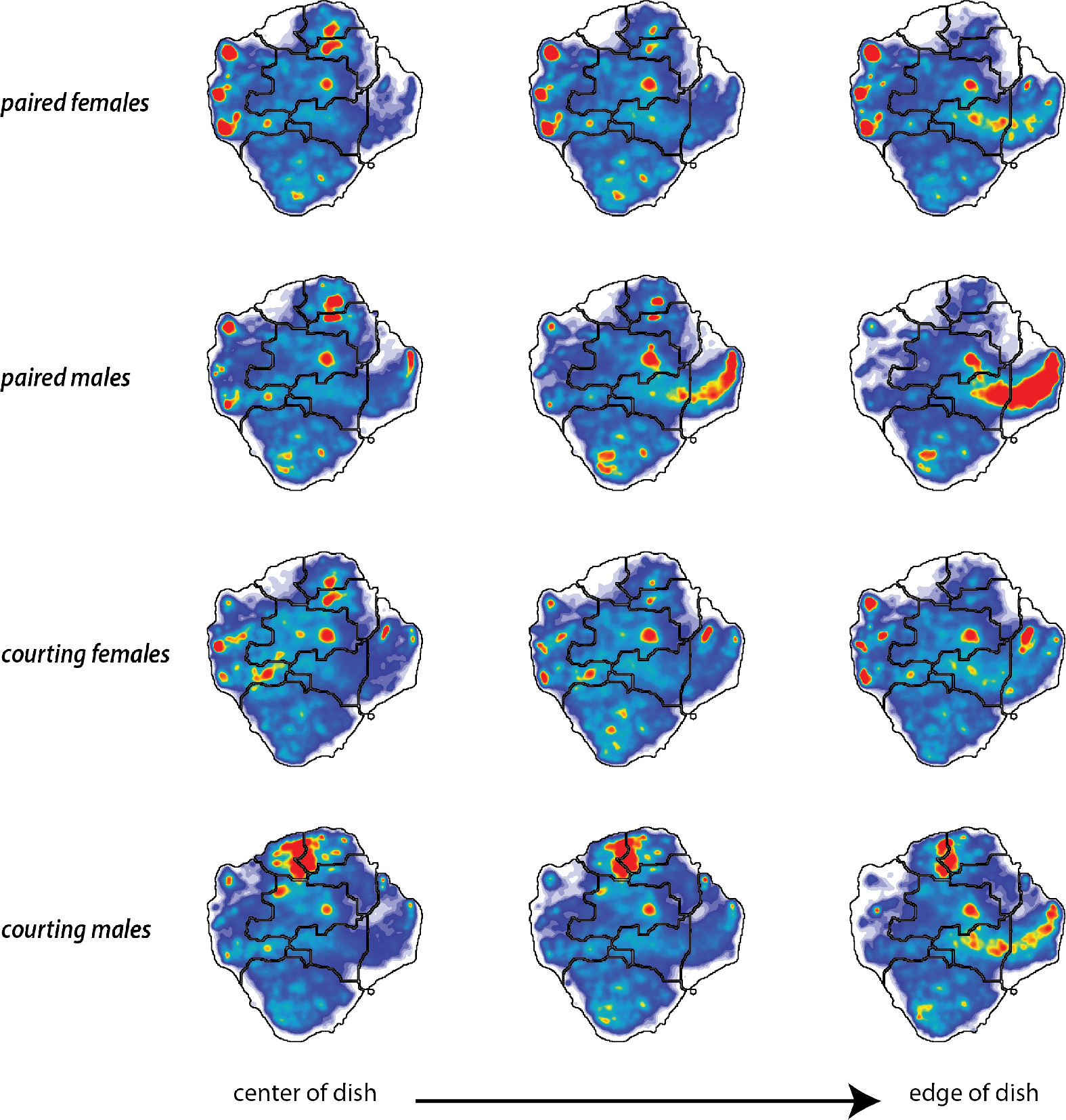
Behavioral density maps of paired individual corresponding to the location in the arena. Density maps are again generated by embedding behaviors exhibited by paired flies while residing in three equal-area concentric circles (centered at R=3.15, 7.66, 9.99mm where *R*_arena_ = 11*mm*) regardless of where the partner is located.

**Figure 5.**
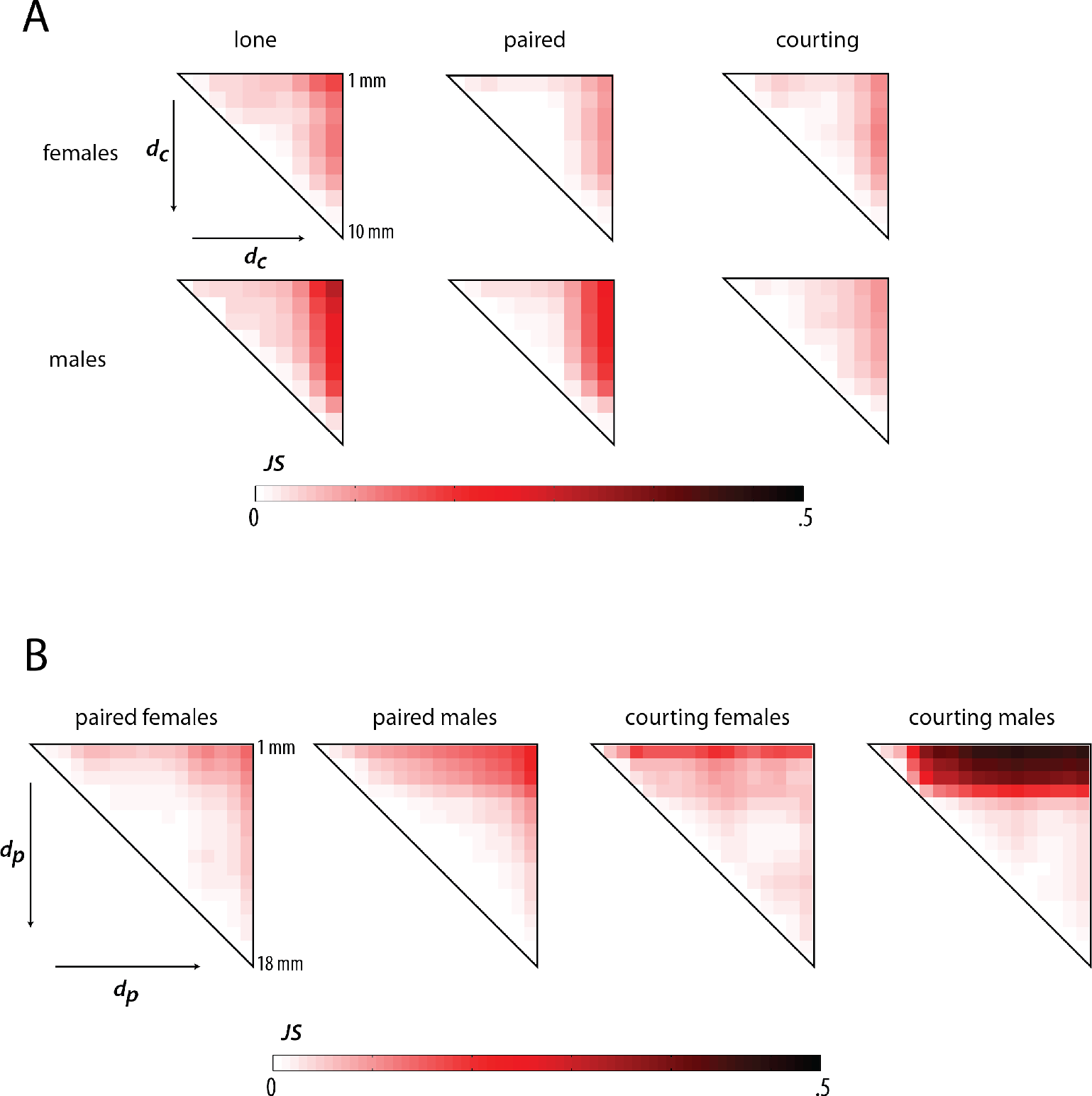
a) Comparison of density maps generated for each context given 1mm bins of distance to the center of the arena. A behavioral map is generated for each context given the behavior of individuals at a specific 1mm window of distance to the center of the arena. The JS divergence is computed between each map within a context, and the upper right section of the resulting matrix is displayed. b) Comparison of density maps for overlapping 2mm bins of distance to partner is generated in the same manner as in part (a) and the JS divergence between each map in a given context is displayed.

### The effect of a partner on behavior

We have previously shown that distance to a courtship partner has an effect on behavioral density (*Klibaite et al., 2017*). We generate behavioral maps for each paired context from time points spend in sliding 2*mm* distance bins to the partner, and find that this is also true for same-sex pairings. We perform the same analysis of divergence between maps as when investigating spatial effects, and discover a blocky structure in the all-to-all JS divergence measurements across partner distance in certain contexts (Fig. 5 B). There appear to be two modes in courting male behavior depending on proximity to the female, and the drop off between these modes occurs at a distance of approximately 4*mm* to the female. Surprisingly the male-male context has a similar behavioral switch at this distance, although the effects are not as strong based on divergence to maps at a greater partner-distance. The changes to the female behavioral density as a function of partner distance are more subtle in either female context, and suggest that social behavior in females is not as dependent on these simple variables.

In order to build towards models of social behavior we must consider whether the individual preferences of animals within a pairing affect the behavior of both individuals within a pairing. We find the correlation values of pairs of coarse-grained behaviors in single individuals and individuals within paired contexts (Fig 6A). There are several consistent trends across all individuals, such as positive correlations between crawling and locomotion, which simply means that individuals that tend to walk often also tend to run more than their counterparts. Another similar finding is that individuals that perform wing-related behaviors often also perform more posterior grooming, likely because these behaviors are related and likely to be performed one after the other based on the hierarchical transition structure we know is a feature of fly behavior (*Berman et al., 2016*). Other features of both lone and paired individuals include negative correlations between vastly different behaviors; for example, there is a fairly strong negative correlation in all cases between crawling and idle behaviors, which means that flies that spend their time running will spend less time standing still. The correlation analysis in individuals simply relates how individuals distribute their time spent performing different behaviors.

**Figure 6.**
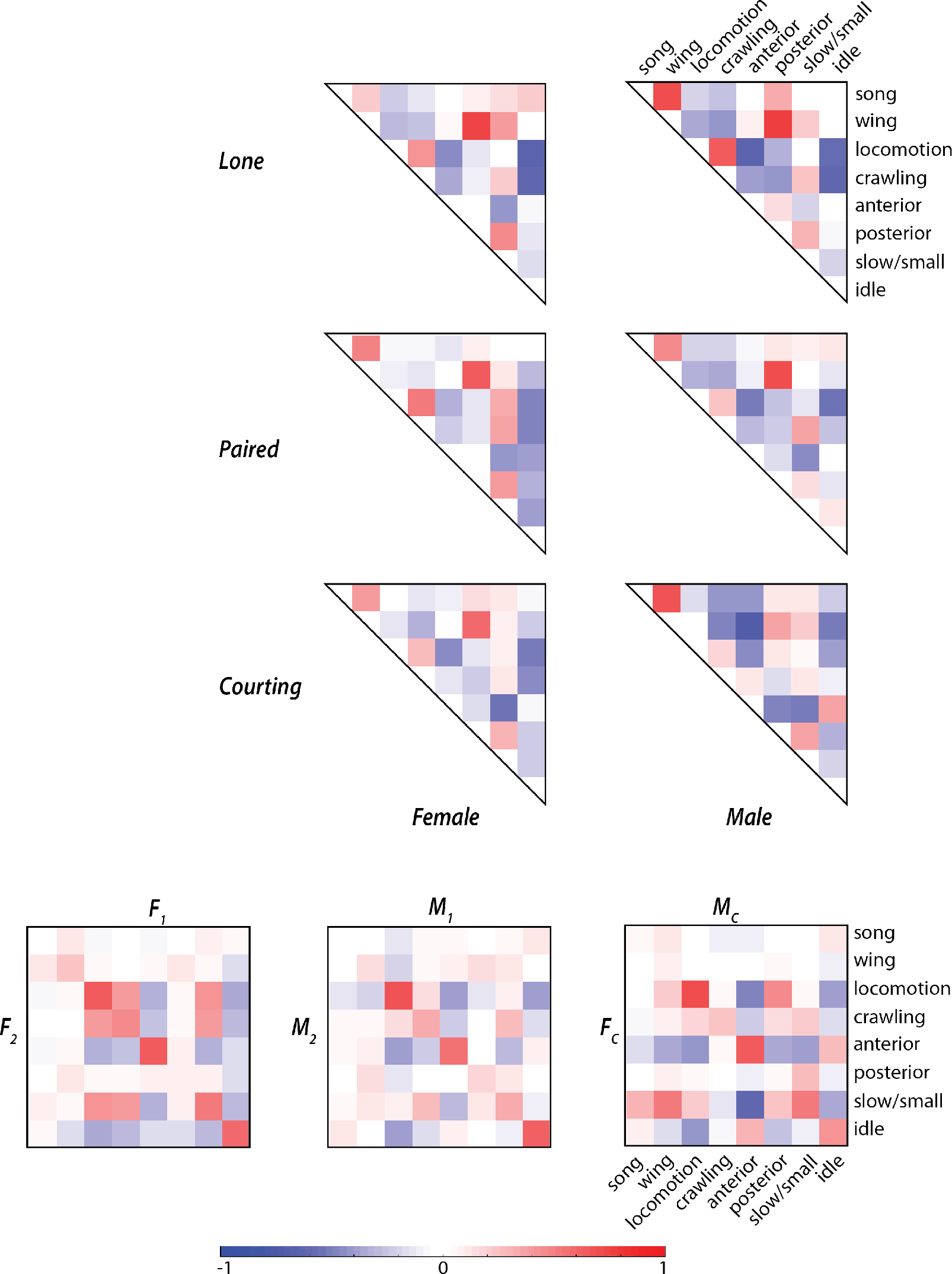
a) The behavioral density given eight coarse behaviors is calculated for each individual in a given context, and the correlation coefficient between all pairs of behaviors over all individuals in specified context is displayed. b) The behavioral density for each individual in paired contexts in calculated as in part (a). For each possible pair of behaviors, the correlation coefficient between behavioral pairs between paired individuals is displayed.

Correlations in behaviors between paired individuals illustrate how a social pairing influences the behavior of each present individual (Fig. 6B). We display the correlation values between coarse-grained behavioral densities of paired individuals and find that there are strong positive correlations across several behavioral pairings. We find that locomotion, anterior grooming, and idle behaviors are moderately correlated in same-sex pairs of flies (correlation values between paired females are .64, .63, and .59, respectively, and correlation values between paired males are .68, .55, and .60, respectively). Courting flies also have a moderate to strong correlation for locomotion (*r* = .69) and anterior grooming (*r* = .65) within pairs, as well as other weaker off-diagonal correlations that are due to courtship specific behaviors such as male wing behaviors and slow female movements (*r* = .52).

Correlation analysis shows that flies will align their behavior when introduced to an arena together, however this tells us nothing about the temporal structure of paired behavior. In order to address whether individuals change their behavior not just due to the proximity of the interaction partner but its behavior we look for structure in the simultaneous behavior of paired flies. From simply watching behavioral movies, we get a sense that sometimes the paired flies are aware of each other in some way, for example by running around the arena together or tapping each other, and that at other times the flies seem totally independent from one another. In order to determine if individuals are interacting instead of simply performing behaviors from an altered distribution in response to the presence of another individual, we consider simultaneous behaviors of paired individuals.

### Synchronization of instantaneous behaviors

The top row of Figure 7 visualizes the expected distribution of simultaneous behaviors given the assumption of independence, whereas the bottom row shows the real distribution of simultaneous behaviors where all individuals have been pooled within each context. We calculate the mutual information present in the simultaneous behavior of paired individuals from this pooled data and display the positive components that contribute to the mutual information in Figure 8A. We find that in all three types of pairs there is a clear enrichment of information along the diagonal, which indicates that paired individuals perform the same behavior more often than is expected by underlying probability alone. This effect comes mainly from simultaneous locomotion, anterior grooming, and idle states, and varies according to pairing. Female-female pairs are particularly likely to remain idle together whereas male-male and courting pairs are more likely to locomote together.

**Figure 7.**
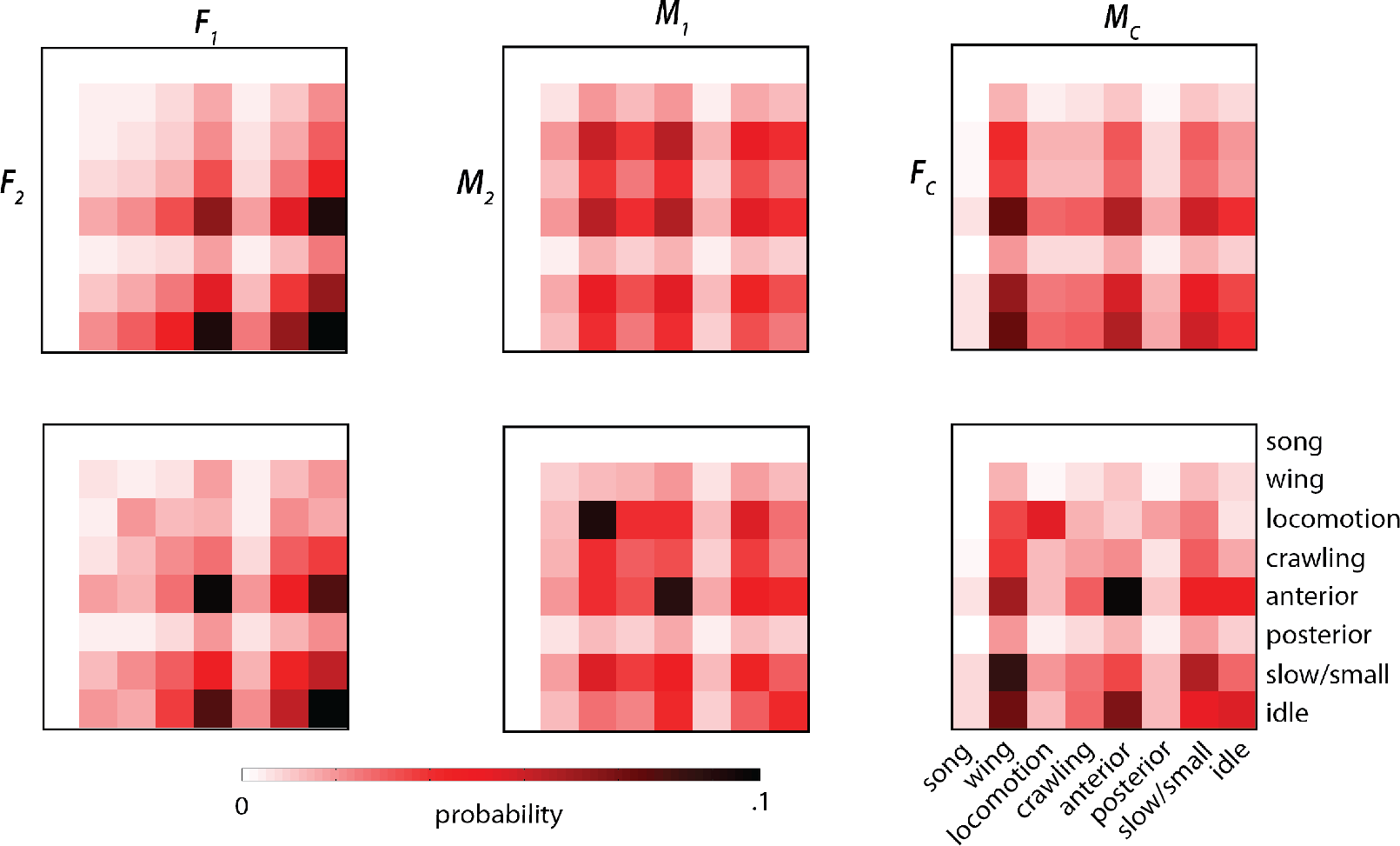
a) The expected joint distribution is found by assuming independence between behaviors performed simultaneously in a given pairing. b) The real joint distribution of behaviors performed simultaneously between individuals in a given context.

**Figure 8.**
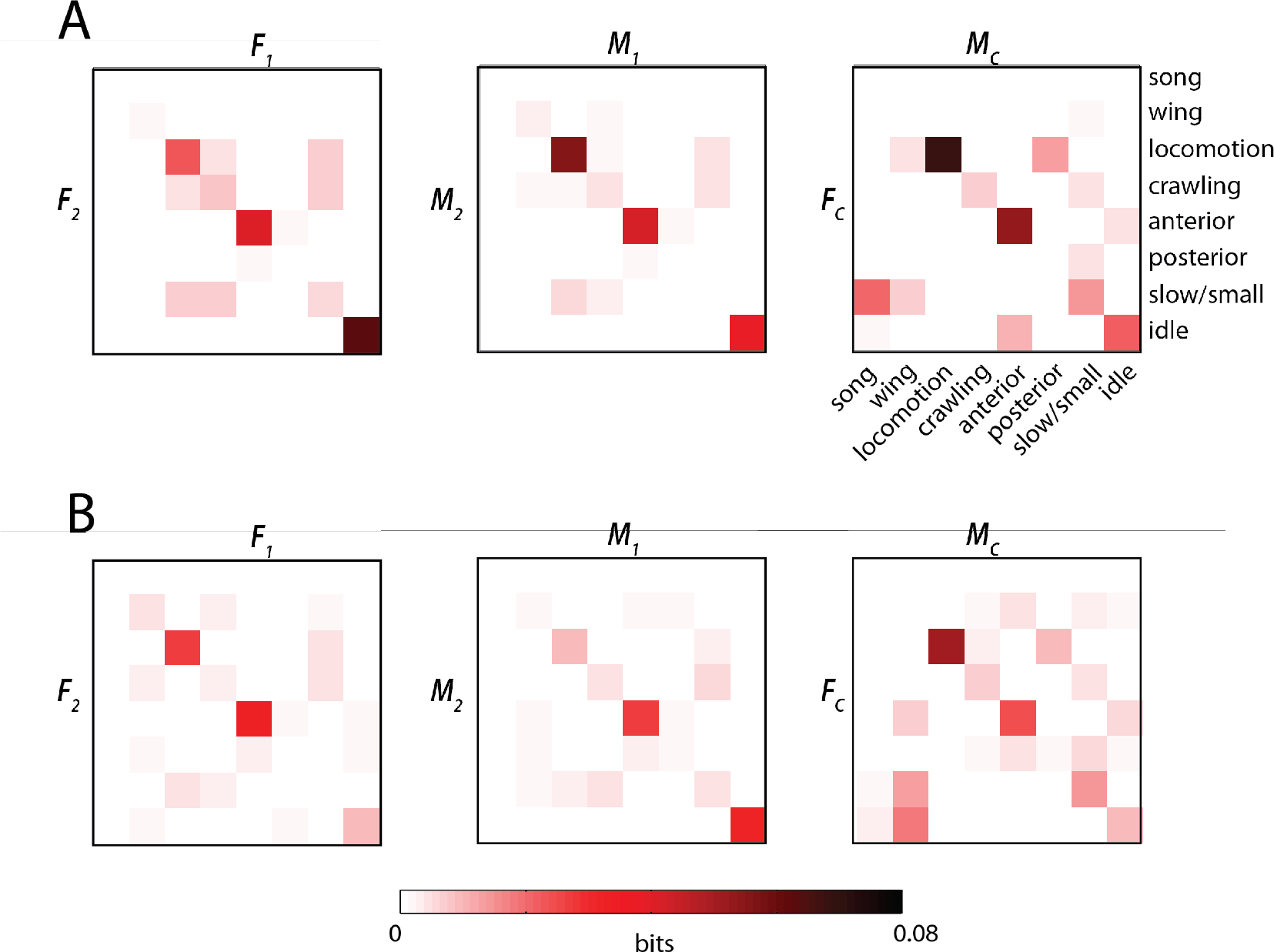
a) The partial mutual information for all data from each behavioral pairing in a paired context is calculated and positive values are displayed. b) The partial mutual information is calculated as in part (a) independently for each pairing in a given context, and the mean value is shown for each set of paired coarse behaviors. Only positive values are displayed

The mutual information in the same-sex pairings is symmetric because each interaction is counted twice, once for each individual. The courting matrix considers the male and female behaviors separately. The non-negative partial mutual information in the first few columns informs us that we have some information about what the female is likely to be doing while the male is singing and performing wing motions, and this is most often idle behaviors, slow movements, and anterior grooming. When we sum over the partial mutual information values in each category, we can calculate the total mutual information for a particular context, summarized in Table 1.

**Table 1.** The sums across partial mutual information are calculated for each context from the matrices displayed in Figures 8A, 8B and from synthetic data generated using a one-step HMM with probability densities and transition data derived from behavioral sequences for each condition. All values are in bits.

| Context | Total MI | Movie-Specific MI | Synthetic Data MI |
| --- | --- | --- | --- |
| Female Pairs | .0954 | .0508 | 5.54E-4 |
| Male Pairs | .0895 | .0488 | 7.67E-4 |
| Courting Pairs | .212 | .110 | 9.15E-4 |

When we calculate the mutual information (MI) on a movie-by-movie basis (Fig. 8B), and consider only the underlying probability for each individual instead of all individuals in that context, we find that the information content is somewhat diminished across each of the contexts. This indicates that paired individuals have some mutual behavioral effect that is not simply related to their simultaneous actions. The enrichment in the simultaneous locomotion and simultaneous anterior entries informs us that even within a pairing these behaviors are more likely to be performed together than by chance. The reduction in paired male simultaneous locomotion and paired female simultaneous idle states, however, indicates that this effect in the combined MI calculation was due to a synchrony effect on an experimental, and not temporal, scale. This means that two females placed together may both become more idle for the course of a recording, but not necessarily synchronize the time spent idle in the way that occurs with other simultaneous behaviors. The same is true for males particularly prone to locomotion.

Histograms of partial mutual information (PMI) for several synchronized behaviors show that not all pairings contribute to positive non-zero MI values (Fig. 10). Courtship pairings with high PMI values for synchronous locomotion indicate times where males spent a long period of time chasing females, but even this interaction was not seen in a significant fraction of recordings. Similarly, each PMI distribution shows that many pairings, even when pooled data shows an effect, did not individually show synchronization.

**Figure 9.**
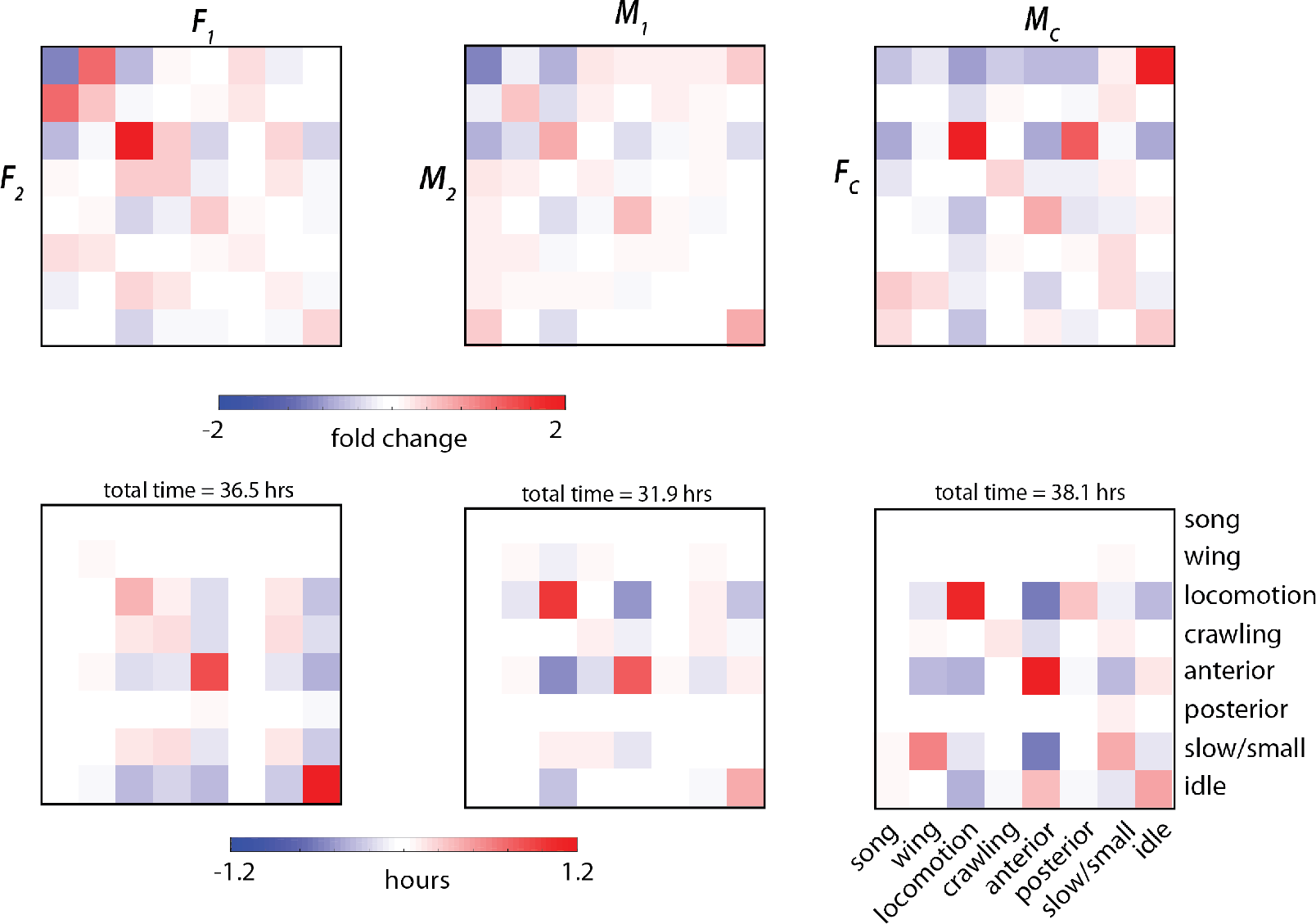
a) The fold change refers to the deviation in fraction of time paired individuals spent performing a set of behaviors simultaneously from the expected probability under the assumption of independence. b) The enrichment in the amount of time spend performing a set of behaviors simultaneously illustrates how much time individuals spent performing a set of simultaneous behaviors above expectation given the combined length of movies in an experiment.

**Figure 10.**
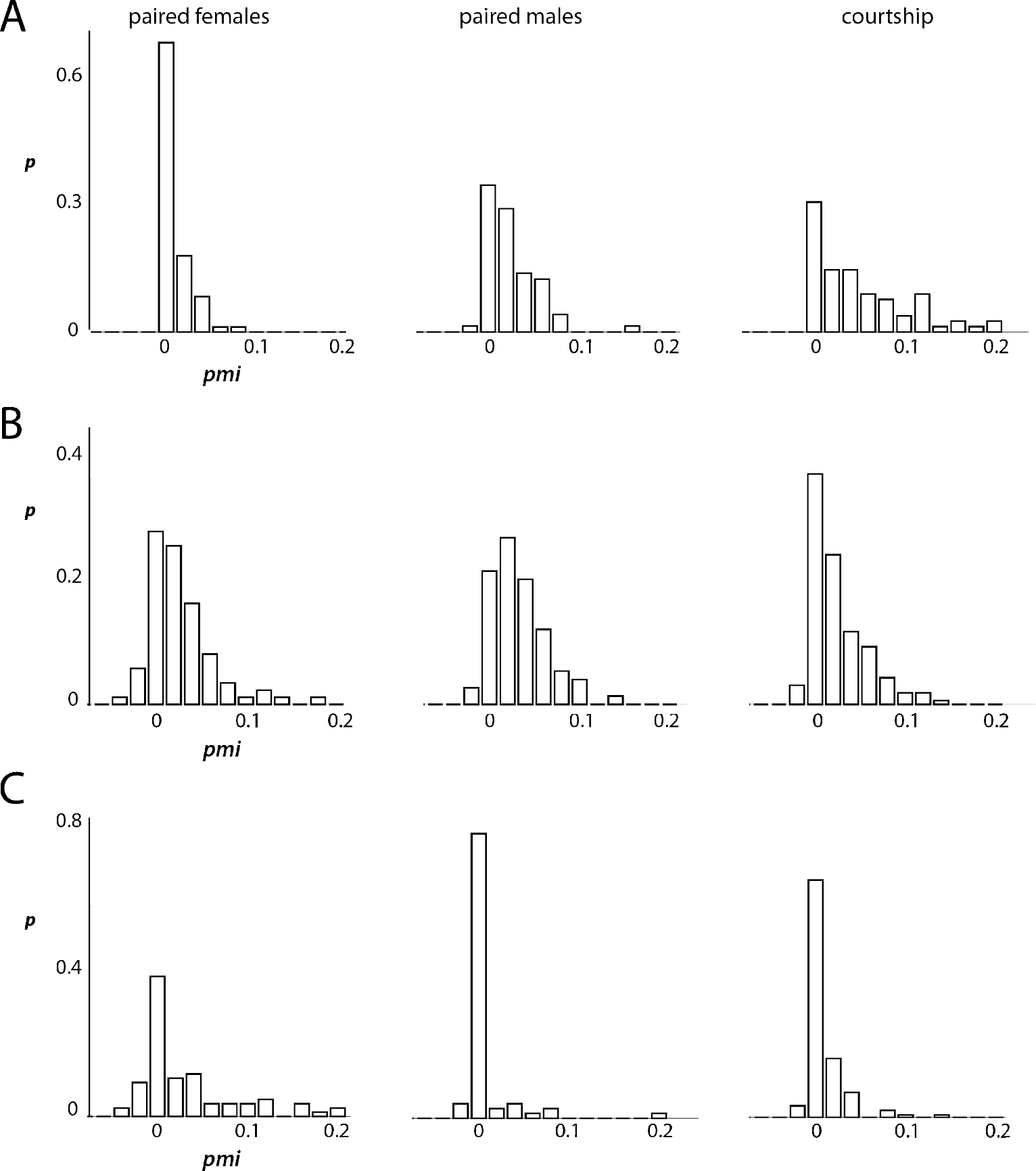
The distribution of partial mutual information values for (a) simultaneous locomotion, (b) simultaneous anterior movements, and (c) simultaneous idle behavior is shown for each of the three paired contexts.

**Figure 11.**
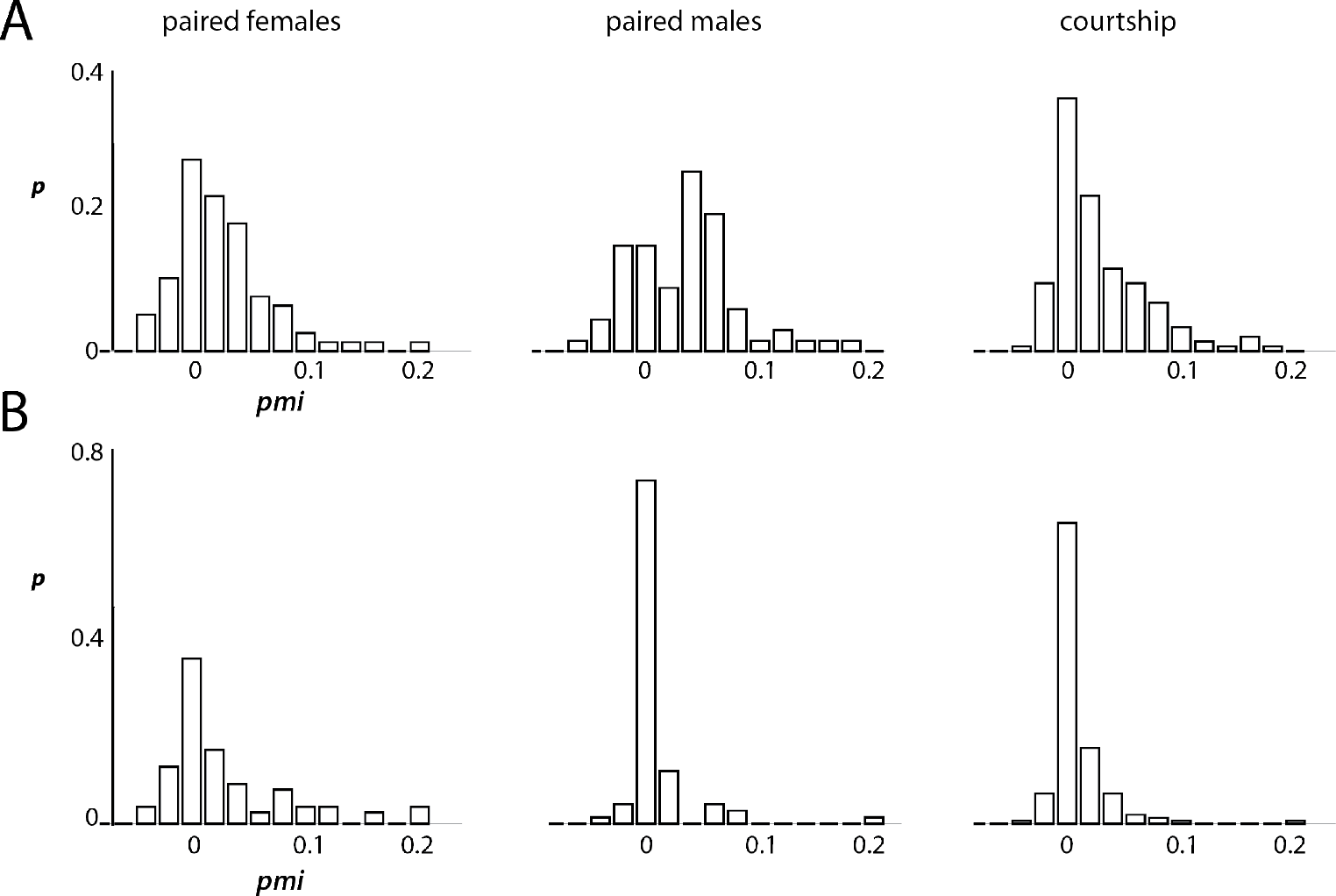
The partial mutual information distributions are calculated across each context for (a) simultaneous anterior movements and (b) simultaneous idle behavior after exclusion of all time points where either of the paired individuals is moving at a velocity above .4*mm*/*s*

One possible explanation for the synchronization we observe based on MI values is that one individual will be startled or begin running based on the activity of the other individual sharing the arena. This could be based on visual cues that do not necessitate the presence of another fly, but instead simply any moving object. After exclusion of time points where either individual in an experiment was moving quickly (threshold = .4*mm*/*s*) we find that the positive distribution of PMIs indicating simultaneous anterior movements is not diminished. This means that even within time spent standing still and performing only small movements, individuals spent more time than expected performing grooming of their heads and antennae than expected by assuming independence.

While there are clear trends for preference of positioning between animals in a small arena (Fig. 1), we also find that the preferential execution of different behaviors depends on an individual’s location in the arena and especially distance to its partner. We visualize these preferences given each of eight coarse behaviors in Figure 12. We observe differences between the behavior of flies between contexts that indicate different aspects of paired behaviors. Females in a courtship condition show few distinctions between behavioral preference, whereas a courting male shows a clear preference for different behaviors based on distance to his courtship target. Courting males differ from males in a same-sex pairing in several interesting ways in particular: they run while near their courtship target (chasing), and perform anterior movements as well as idle behavior more rarely when near the interaction partner. These preferences, coupled with a propensity to perform wing extension and wing motions (song) when near the female result in canonical courtship behavior.

**Figure 12.**
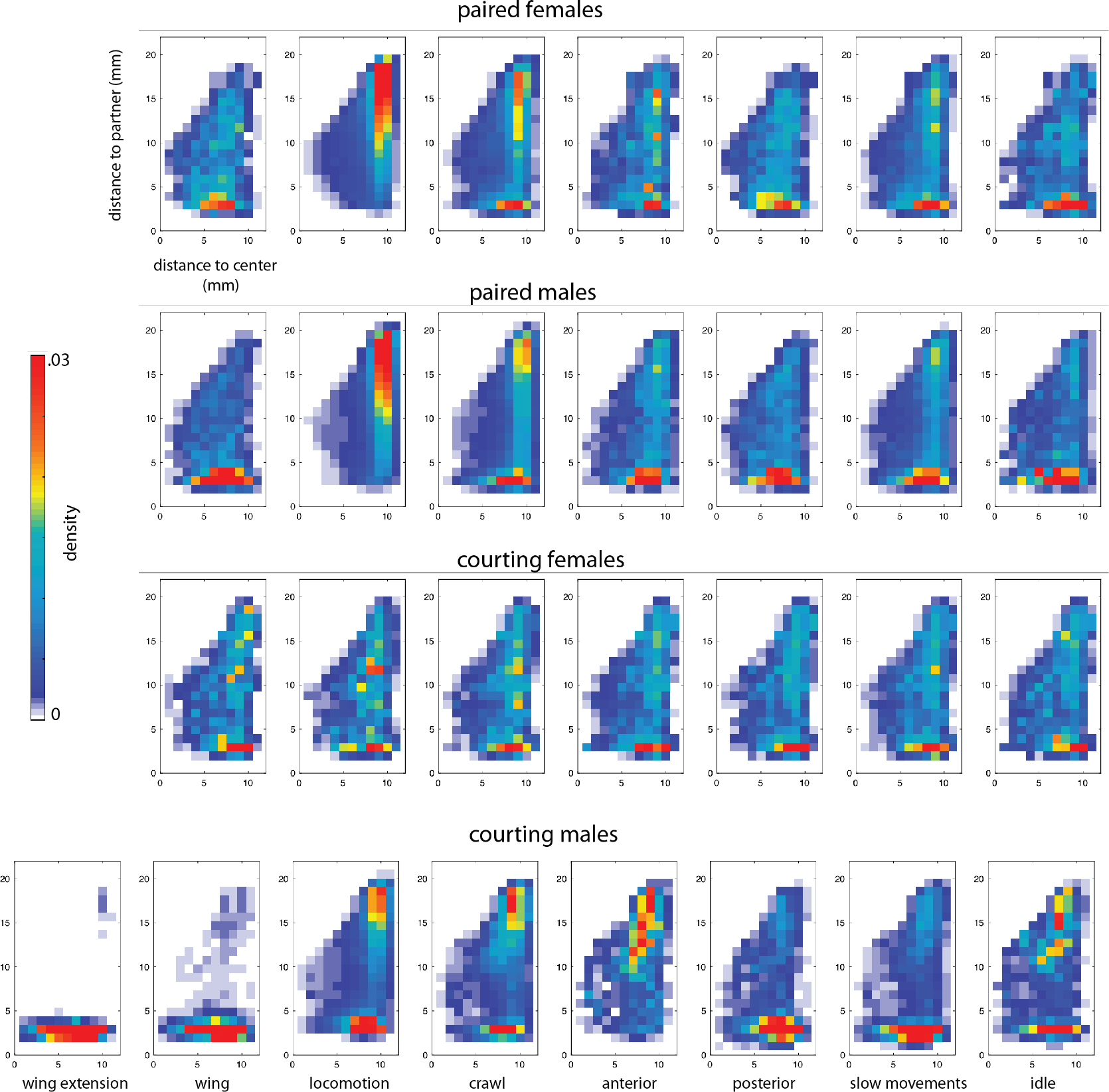
For each of eight behaviors (columns) for each paired context (rows), the 2-dimensional histogram of residence time in each paired context is broken down by distance from the center of the arena (x-axis) and distance to the interaction partner (y-axis). Each heat map summarizes where an individual was in the arena and relation to the interaction partner when performing a particular behavior.

## Discussion

The promising trend at the intersection of traditional ethology and quantitative techniques has crystallized questions about the explanatory power of behavior (*Schaefer and Claridge-Chang, 2012*; *Krause et al., 2013*; *Anderson and Perona, 2014*). A major aim of neuroscience is the untangle the genetic and neural underpinnings of the actions of an animal while acting and interacting with its environment, where we believe we are reading out some noisy downstream output of many sensory and neural transformations. We can ascribe behaviors to causes originating from neural activity and design experiments to test these assumptions. However, we also have deeper questions in regards to the nature of how and why behavior is organized in the brain, in time, and between multiple individuals in order to produce collective effects. There is a newfound, or perhaps, in the face of new high-throughput techniques, seemingly more tangible, desire to ascribe principles to the organization of behavior, and to explain how behaviors have evolved to assume such organization. We refer to the combined effects of these properties as the structure of behavior.

We know many particulars about behaviors of and interaction between fruit flies, such as the averaged responses to particular pheromones or stimuli (*Sokolowski, 2010*; *Agrawal et al., 2014*) but only have a vague sense of what will happen when they are allowed to freely interact in a naturalistic environment. After introducing two conspecifics, how much do we really know about how an interaction will progress? In order to address behavioral dynamics in a tangible way, we ask if there is structure in the behavior of the fruit fly that is present regardless of pairing and if there are special contexts or circumstances that predictably alter this structure.

Here we address the structure of paired behavior in fruit flies at several scales, paying particular attention to the differences between paired and lone flies. We wish to discern the effect that simply being in a social context has on behavior, as well as how animals respond to different contexts within the large umbrella of socialization. We calculate features that we can keep track of throughout our analysis, such as the distance between individuals and the deviation in their heading changes, and show how these shape the density distribution of the behaviors performed by an animal. We address the same effects in lone animals where possible, and show that unpaired animals respond to their geometry, and these effects persist in the paired contexts.

We illustrate several ways in which social behaviors may emerge with a simple example in Figure 13. A toy ethogram and behavioral density bar plot are drawn for a (red) fly in an isolated condition, as well as in several cases when paired with a second (blue) fly. Case I represents the behavior of a hypothetical lone individual that performs three behaviors with equal frequency. Case II represents the case where this individual does not change its behavioral frequency in the presence of a second individual, however the ethograms indicate that individuals have synchronized their behavior. In Case III the individual performs the three behaviors seen in the lone case, but with an altered distribution. Finally, Case IV represents the case where a new behavior is performed in the presence of a social partner.

**Figure 13.**
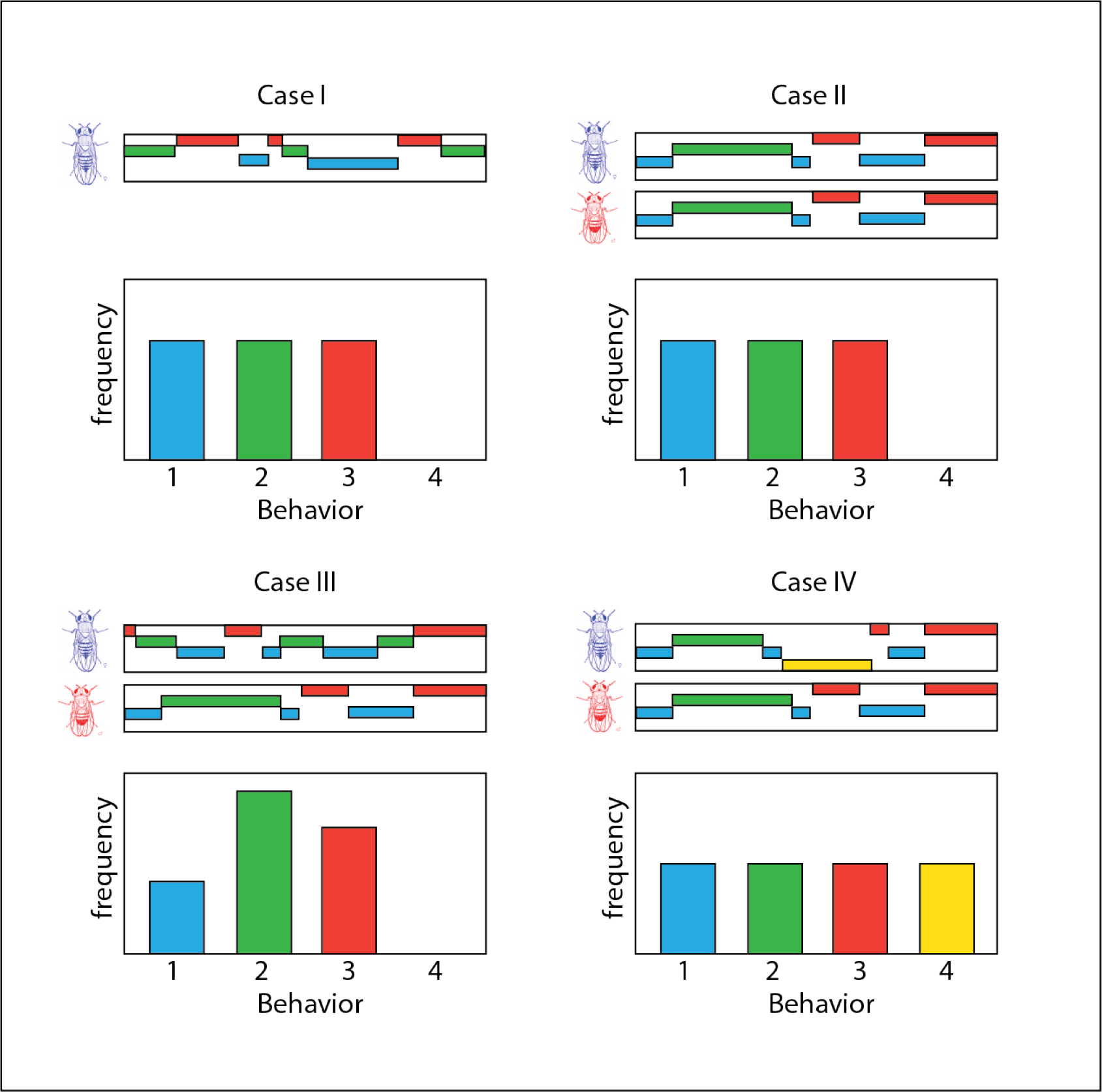
A summary of ways in which individuals may alter behavior during social interaction. Case I shows a toy ethogram and behavioral density of a lone fly. Case II represents a social interaction where behavioral density does not change, but temporal synchrony occurs. Case III represents a change to the behavioral density of the first individual. Case IV represents the introduction of a novel behavior due to interaction.

We see evidence of each of these types of structure in the social interactions of fruit flies. Case IV demonstrates the introduction of a new social behavior, which we observe when courting males initiate song, and is the most apparent shift from lone to social behavior. The use of unsupervised techniques that can produce behavioral labels for large amounts of data have allowed identification of other more subtle types of interactions. We see a shift in behavioral density not only when pooling data over many individuals (Fig. 2) but also within individuals experiencing different social situations (Fig. 6). The correlation coefficients of certain behaviors (namely anterior-anterior, locomotion-locomotion, and idle-idle) between pairings of individuals within a given pairing indicate that each paired set of flies is unique but on average pairs tend to adjust or match their behaviors throughout an interaction. Mutual information analysis on pooled and individual interaction experiments confirms this matching, as well as indicates an even more precise temporal component of behavioral synchronization. Even by excluding data points where either individual was moving within the experimental chamber, we find that individuals perform anterior movements together more often than expected when assuming independence.

These aspects of behavioral organization offer explanation for how more complex, and potentially collective, social behaviors are constructed. Groups of flies may use synchronization in social situations to successfully navigate new environments by leveraging information from their conspecifics. Some social imperatives, such as that of the male to chase and sing to the female, may outweigh these subtle effects. One aspect of the synchronization we observe is that it is inconsistent across pairs even within the same context. The stochastic nature of behavior may balance organizational principles and lend to exploration and variation.

## Methods and Materials

### Behavioral Movie Recordings

We use a custom-built rig to film the behavior of interacting fruit flies. Our rig can accommodate four experiments at once, allowing for the acquisition of two dozen half-hour movies per day. The setup is the same one described in (*Klibaite et al., 2017*), where four cameras are used at once to record the activity in four separate domes placed on the same backlight. Flies are loaded by gentle manual aspiration and movies are started within a minute after flies are introduced into the arena. Because fruit fly activity and courtship levels are known to vary over their circadian cycles and lifetimes, we aim to capture behavior from flies at similar times within these cycles and film only within the four hours after lights come on. To keep experiments consistent, we isolate flies upon eclosion and age them four to six days before imaging.

In order to address the effects of paired social context, and not just the courtship context, we filmed behavioral movies of male-female pairs as well as male and female same-sex pairs. Additionally, movies of isolated flies from the same population for either sex provide a control that allows us to compare spontaneous behavior to interaction. We have increased the time captured for each type of interaction from what was previously described in order to achieve high enough sampling to analyze not only behavioral densities but to obtain many samples of each behavior performed. The number of movies and time of behavior recorded for each pairing is summarized in Table 2. Courtship movies are not analyzed past the point of successful copulation, and therefore have variables lengths, whereas all other movies are analyzed over the entire 30 minutes of recording.

**Table 2.** Number of trials for each behavioral context and total recording time from movies analyzed are listed for each context.

| Social Context | Number of Movies | Behavioral Hours |
| --- | --- | --- |
| Courtship (Female) | 152 | 38.1 |
| Courtship (Male) | 152 | 38.1 |
| Female-Female | 81 | 73.1 |
| Male-Male | 70 | 63.9 |
| Lone Female | 61 | 30.5 |
| Lone Male | 55 | 27.5 |

### Behavioral Embeddings

We perform behavioral embeddings for all recorded movies using the pipeline introduced in (*Berman et al., 2014*). Table 3 lists the parameters used for image PCA, wavelet decompositions, tSNE embedding, and clustering. The original embedding was performed using important sampling from all movies in order to facilitate comparisons between the various social contexts. after embedding we sampled movies from each cluster of the map after performing the watershed transform and attached descriptive terms in order to combine clusters into coarse grained regions.

**Table 3.** Parameters used in the behavioral analysis pipeline are described and values used throughout this analysis are listed. For more details see supplement of (*Berman et al., 2014*)

| Parameter | Value | Description |
| --- | --- | --- |
| $N_{\theta}$ | 90 | Number of angles used in Radon transform |
| $M$ | 50 | Number of postural eigenmodes used for projections |
| $N_f$ | 25 | Number of frequency channels used in wavelet decomposition |
| $N_{train}$ | 38,500 | Training set size after subsampling |
| $\omega$ | 1.5 | Width of Gaussian kernel used for generating density map |

### Density Map Distance Metrics

The distance between embeddings between different contexts as seen in Figures 2 and 5 is the Jenson-Shannon divergence as defined in Equation 1 where the Kullback-Leibler divergence (KL) is defined as in Equation 2 (*Lin, 1991*). Since any behavioral map (represented by a 501×501 matrix) may have very small values, a mask is first applied to each map before comparison so that only entries with value *>* 1*E* − 6 in the mean behavioral map are used. All infinite and NaN values are then removed before summation. Units are presented in bits.

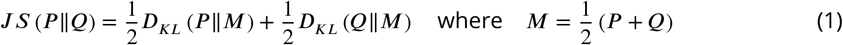

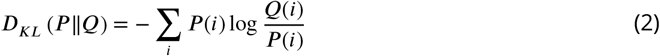

### Calculating Behavior Correlations

Densities for each of eight coarse behaviors are determined by the fraction of time an individual spent in those states as determined by embedding in the behavioral map. All densities sum to one. Correlations between behavioral probabilities within individuals and between paired individuals are measured using the Pearson’s linear correlation coefficient.

### Mutual Information Analysis

We calculate the mutual information for all movies in a given context, as well as each movie individually. We use with the following equation, where *B*_1_ and *B*_2_ are the underlying distributions of behaviors for a given type of individual.

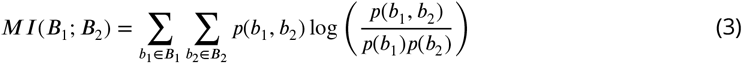

We produce synthetic data given an 8-state HMM (Hidden Markov Model) with the probability and transition matrices calculated from trajectories through the coarse behavioral space for each interaction context. The synthetic data is the same length as the true data for each context.

## Acknowledgments

The authors acknowledge support by the National Science Foundation, through the Center for the Physics of Biological Function (PHY-1734030) and award IOS-1451197, and the National Institutes of Health (R01NS104899). This work was performed in part at the Aspen Center for Physics, which is supported by National Science Foundation grant PHY-1607611.

## References

Agrawal S, Safarik S, Dickinson M. The relative roles of vision and chemosensation in mate recognition of Drosophila melanogaster. Journal of Experimental Biology. 2014; 217(15):2796–2805.

Allee WC, et al. Animal aggregations. 1931;.

Anderson DJ, Perona P. Toward a science of computational ethology. Neuron. 2014; 84(1):18–31.

Bates LA, Byrne RW. Imitation: what animal imitation tells us about animal cognition. Wiley Interdisciplinary Reviews: Cognitive Science. 2010; 1(5):685–695.

Berman GJ, Bialek W, Shaevitz JW. Predictability and hierarchy in Drosophila behavior. Proceedings of the National Academy of Sciences. 2016; 113(42):11943–11948.

Berman GJ, Choi DM, Bialek W, Shaevitz JW. Mapping the stereotyped behaviour of freely moving fruit flies. Journal of The Royal Society Interface. 2014; 11(99):20140672.

Bialek W, Cavagna A, Giardina I, Mora T, Pohl O, Silvestri E, Viale M, Walczak AM. Social interactions dominate speed control in poising natural flocks near criticality. Proceedings of the National Academy of Sciences. 2014; 111(20):7212–7217.

Byrne RW. Animal imitation. Current Biology. 2009; 19(3):R111–R114.

Censi A, Straw AD, Sayaman RW, Murray RM, Dickinson MH. Discriminating external and internal causes for heading changes in freely flying Drosophila. PLoS computational biology. 2013; 9(2):e1002891.

Coen P, Murthy M. Singing on the fly: sensorimotor integration and acoustic communication in Drosophila. Current opinion in neurobiology. 2016; 38:38–45.

Coen P, Xie M, Clemens J, Murthy M. Sensorimotor transformations underlying variability in song intensity during Drosophila courtship. Neuron. 2016; 89(3):629–644.

Dombrovski M, Poussard L, Moalem K, Kmecova L, Hogan N, Schott E, Vaccari A, Acton S, Condron B. Cooperative Behavior Emerges among Drosophila Larvae. Current Biology. 2017; 27(18):2821–2826.

Durisko Z, Kemp R, Mubasher R, Dukas R. Dynamics of social behavior in fruit fly larvae. PLoS One. 2014; 9(4):e95495.

Giuggioli L, Potts JR, Rubenstein DI, Levin SA. Stigmergy, collective actions, and animal social spacing. Proceedings of the National Academy of Sciences. 2013; 110(42):16904–16909.

Herbert-Read JE, Perna A, Mann RP, Schaerf TM, Sumpter DJ, Ward A J. Inferring the rules of interaction of shoaling fish. Proceedings of the National Academy of Sciences. 2011; 108(46):18726–18731.

Iacoboni M. Imitation, empathy, and mirror neurons. Annual review of psychology. 2009; 60:653–670.

Katz Y, Tunstrøm K, Ioannou CC, Huepe C, Couzin ID. Inferring the structure and dynamics of interactions in schooling fish. Proceedings of the National Academy of Sciences. 2011; 108(46):18720–18725.

Klibaite U, Berman GJ, Cande J, Stern DL, Shaevitz JW. An unsupervised method for quantifying the behavior of paired animals. Physical biology. 2017; 14(1):015006.

Krause J, Krause S, Arlinghaus R, Psorakis I, Roberts S, Rutz C. Reality mining of animal social systems. Trends in ecology & evolution. 2013; 28(9):541–551.

Lin J. Divergence measures based on the Shannon entropy. IEEE Transactions on Information theory. 1991; 37(1):145–151.

Lorenz K. On aggression. Psychology Press; 2002.

Louis M, de Polavieja G. Collective Behavior: Social Digging in Drosophila Larvae. Current Biology. 2017; 27(18):R1010–R1012.

Ni R, Ouellette N. Velocity correlations in laboratory insect swarms. The European Physical Journal Special Topics. 2015; 224(17-18):3271–3277.

Ramdya P, Schneider J, Levine JD. The neurogenetics of group behavior in Drosophila melanogaster. Journal of Experimental Biology. 2017; 220(1):35–41.

Rizzolatti G, Craighero L. The mirror-neuron system. Annu Rev Neurosci. 2004; 27:169–192.

Schaefer AT, Claridge-Chang A. The surveillance state of behavioral automation. Current opinion in neurobiology. 2012; 22(1):170–176.

Shorey H, Bartell R. Role of a volatile female sex pheromone in stimulating male courtship behaviour in Drosophila melanogaster. Animal behaviour. 1970; 18:159–164.

Sokolowski MB. Social interactions in “simple” model systems. Neuron. 2010; 65(6):780–794.

Stowers JR, Hofbauer M, Bastien R, Griessner J, Higgins P, Farooqui S, Fischer RM, Nowikovsky K, Haubensak W, Couzin ID, et al. Virtual reality for freely moving animals. Nature methods. 2017; 14(10):995.

Tinbergen N. On aims and methods of ethology. Ethology. 1963; 20(4):410–433.

Tolman CW. Social facilitation of feeding behaviour in the domestic chick. Animal Behaviour. 1964; 12(2-3):245–251.

Valente D, Golani I, Mitra PP. Analysis of the trajectory of Drosophila melanogaster in a circular open field arena. PloS one. 2007; 2(10):e1083.

Weisstein EW. Disk line picking. 2000;.

Welty JC. Experiments in group behavior of fishes. Physiological Zoology. 1934; 7(1):85–128.

Zentall TR. Action imitation in birds. Animal Learning & Behavior. 2004; 32(1):15–23.

Zentall TR. Imitation: definitions, evidence, and mechanisms. Animal cognition. 2006; 9(4):335–353.

